# Attentive red squirrel mothers have faster-growing pups and higher lifetime reproductive success

**DOI:** 10.1101/594093

**Authors:** Sarah E Westrick, Ryan W Taylor, Stan Boutin, Jeffrey E Lane, Andrew G McAdam, Ben Dantzer

**Author notes:** Corresponding author, 530 Church St., Ann Arbor, MI 48109.

## Abstract

Parental investment theory predicts that observed levels of parental care afforded to offspring are set by the benefits (to offspring quality and survival) relative to the costs (to parental survival or future reproduction). Although difficult to document in mammals, there is often substantial individual-variation in the amount of parental care within species. We measured the impact of individual variation in maternal care (“attentiveness” towards offspring or maternal motivation) on offspring growth and survival in a wild population of North American red squirrels (*Tamiasciurus hudsonicus*). We used latency to return to pups following a nest intrusion as a measure of maternal attentiveness to pups. We found this behavior to be repeatable within individuals suggesting this behavior is a personality trait or a “maternal style”. In this population, postnatal growth rate is important for pup overwinter survival. Pups from large litters grew faster if they had a highly attentive mother, indicating that maternal care behavior can mitigate the trade-off between litter size and offspring growth and potentially improve survival of pups. Additionally, more attentive mothers had slightly higher lifetime reproductive success than less attentive mothers. These results highlight important fitness effects of having a highly attentive mother and show that maternal care behavior can alter a fundamental life history trade-off between offspring quantity and quality.

**Lay Summary:** It pays to be attentive to your pups as a squirrel mom. In a long-term study of a wild population of North American red squirrels, we observed repeatable individual variation in maternal attentiveness towards offspring. Mothers who returned faster to pups following a nest intrusion produced faster growing pups and were able to produce larger fast-growing litters. Over their entire lifetime, attentive mothers also had more offspring recruit into the breeding population.

## Introduction

Theory predicts that elevated parental investment will produce more, and/or higher quality, offspring, but this may come at a cost of decreased future fecundity and/or survival of parents (Trivers 1974; Drent and Daan 1980; Stearns 1989; Clutton-Brock 1991). In this context, parental care has been defined to include parental traits, including behavior, that increase fitness of offspring, and is one aspect of parental investment (Clutton-Brock 1991; Royle et al. 2012). The energetic costs of reproduction, which may limit future parental investment (Drent and Daan 1980), may be especially high in mammals as offspring are highly dependent on their mother for survival until weaning (Roff 1998; Reinhold 2002; Maestripieri and Mateo 2009). Under the assumption of fixed energy budgets, these energetic costs to parents of parental care likely contribute to the fundamental life history trade-off between offspring number and size (Smith and Fretwell 1974; Charnov and Ernest 2006). This trade-off is reflected in the frequent observation that parents can rarely raise many large offspring (Rogowitz 1996), except when they have a large amount of resources to do so (van Noordwijk and de Jong 1986; Reznick et al. 2000). The trade-off for parents between offspring quality and quantity could impact the lifetime trajectory of offspring via differences in developmental rates of individuals (Clutton-Brock 1991; Royle et al. 2012; Klug and Bonsall 2014). For example, offspring growing up in large litters may exhibit slower development and be smaller at the age of independence than those from smaller litters which could in turn influence lifetime reproductive success (Lindström 1999).

In mammals, we might expect to see directional positive selection for high maternal care if the benefits afforded to offspring exceed the costs to the mother, yet there is often much individual variation in this behavior (Bales et al. 2002; Champagne et al. 2007; Fairbanks and Ackles 2012). For example, in primates, individual variation in maternal care, or “maternal style”, can include protective, rejecting, restrictive, or laissez-faire mothers (Fairbanks and Ackles 2012). Maternal styles can have important consequences for a female’s fitness and may be a mechanism for transmitting individual differences across generations (Fairbanks and Ackles 2012). Many studies in mammals have investigated the proximate mechanisms behind individual variation in maternal care (Numan and Insel 2006), yet the ultimate mechanisms (i.e. fitness consequences) are rarely addressed, especially in wild rodents, due to the logistical challenges of accurately documenting parental behaviors. Much research in primates has been done to describe individual variation in maternal care (Fairbanks and Ackles 2012), however it is challenging to follow the fitness consequences beyond interbirth interval in such long-lived species.

In this study, we used latency to return to pups, a measurement commonly used with laboratory rodents (e.g. Numan and Insel 2006; Champagne et al. 2007), as a measurement of maternal attentiveness following a nest intrusion in wild North American red squirrels (*Tamiasciurus hudsonicus*). In laboratory studies of rodents, latency to retrieve pups after they are moved to a different location in the cage is interpreted as how motivated an individual is to attend to pups, whether they are their own offspring or another’s (Mann 1993; Olazábal et al. 2013). For example, virgin female laboratory rats can develop maternal motivation to retrieve pups by prolonged exposure to pups (Seip and Morrell 2008). Many neuroendocrine studies measure retrieval behavior when manipulating brain regions to identify neural structures involved in maternal behavior (Numan and Woodside 2010). In wild animals, we cannot distinguish between whether this is how motivated a mother is to defend pups or how motivated she is to exhibit infant-directed behaviors, both important components of motivated maternal care (Olazábal et al. 2013).

We conducted this study as part of a long-term project on red squirrels in Yukon, Canada. Red squirrel pups are altricial and dependent upon their mother until weaning whereupon they typically disperse to nearby territories (mean = 92 m – 102 m from natal territory (Berteaux and Boutin 2000; Cooper et al. 2017)). Weaned pups experience strong selective pressures with a majority of juvenile mortality (68%, on average though this is highly variable among years) occurring over the summer before mid-August and few surviving through their first winter (61% of those alive in August, on average), depending on food availability and adult density (McAdam et al. 2007; Hendrix et al. 2019). Pups that grow faster early in life are typically more likely to survive their first winter (McAdam and Boutin 2003a; Fisher et al. 2017; Hendrix et al. 2019), particularly in years with high conspecific density (Dantzer et al. 2013). Despite the link between increased survival and fast growth in red squirrels, there is substantial variation in pup growth rates among and within litters. Some of this variation in pup growth rate is explained by large maternal effects including variation in litter size and the birth dates of litters (McAdam et al. 2002) or levels of maternal glucocorticoids during pregnancy (Dantzer et al. 2013), although much variation among females in the growth rates of their pups remains unexplained (McAdam et al. 2002). Here, we tested the hypothesis that maternal behavior affects offspring growth rate and survival. We also tested the within-individual repeatability of this behavior to identify if red squirrels exhibit a “maternal style” and tested whether more attentive mothers had more pups recruit into the breeding population over their lifetime. We predicted that mothers who exhibited higher maternal attentiveness (i.e., faster return to the nest) would produce faster growing offspring and the pups would be more likely to survive to autumn. Consistent with life history theory, individual growth rates of pups are typically lower in larger litters (Humphries and Boutin 2000), however previous research has shown mothers in a high conspecific density environment or those with supplemental food can reduce this trade-off between litter size and pup growth rates (Dantzer et al. 2013). Therefore, we also hypothesized that the attentiveness of mothers towards their offspring could be one mechanism by which mothers ameliorate this life history trade-off.

## Methods

### Study population

North American red squirrels are arboreal, solitary, and sexually monomorphic (Boutin and Larsen 1993). Both sexes defend individual year-round territories and have many nests on their territory (Dantzer et al. 2012; Siracusa et al. 2017). Mothers provide all parental care in this species and typically produce one successful litter per year, with the exception of white spruce (*Picea glauca*) mast years when they may produce 2 litters (Boutin et al. 2006). Renesting following a failed litter also occurs (Williams et al. 2014). Our study was conducted within Champagne and Aishihik First Nations traditional territory (with their explicit agreement) in Yukon, Canada (61°N, 138°W). Squirrels in our study population rely mainly on the seeds of white spruce for food (Fletcher et al. 2010; Fletcher et al. 2013), which they cache in an underground larder hoard (“midden”) within their territory. White spruce exhibit mast seed production, meaning that trees synchronously produce large numbers of spruce cones in some years followed by years with almost no cones produced (LaMontagne and Boutin 2007). Mast events by white spruce typically occur every 4-6 years (Nienstaedt and Zasada 1990). As their main seed predator in the region, the population density of red squirrels increases following mast seed events (Dantzer et al. 2013; McAdam et al. 2019). Because we have previously documented that spruce cone availability impacts growth and survival of offspring (Humphries and Boutin 1996; Boutin et al. 2000; McAdam and Boutin 2003a; Fletcher et al. 2010; Fletcher et al. 2013), we estimated the number of cones available in the study area by using visual cone counts to determine cone index (for details, see LaMontagne et al. 2005).

### Maternal behavior observations

In 2008, 2009, 2016 and 2017, we live-trapped (Tomahawk Live Trap, Tomahawk, WI, USA) breeding females (n = 272 unique squirrels across 4 years) at regular intervals to determine reproductive status (see McAdam et al., 2007 for more details). Squirrels in this study were from either a control study area (n = 141 squirrels) or a study area that was provided with supplemental *ad libitum* peanut butter from 2004 to 2017, resulting in a higher density of squirrels (n = 79 squirrels) (described in Dantzer et al. 2013). Squirrels in the high-density study area typically have higher levels of glucocorticoids (Dantzer et al. 2013) and spend less time in the nest (Dantzer et al. 2012). We included these squirrels from the high-density study area because it increased our sample size by 56%. However, in our present study, study area did not predict growth rate (Table 1). Nonetheless, to control for any variation due to this difference in conspecific density and food availability, we included study area as a covariate in all models predicting pup survival or growth rate.

**Table 1.**
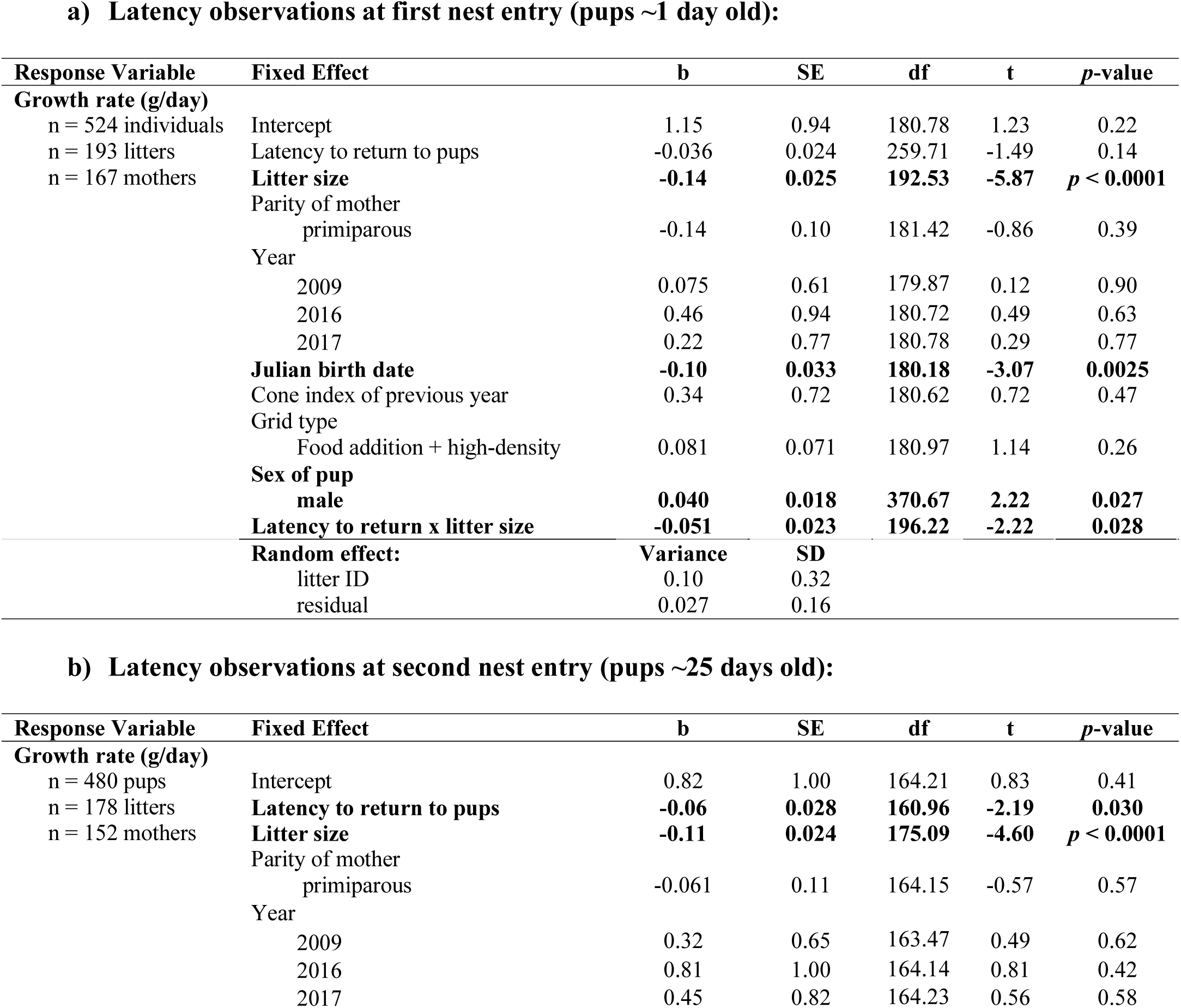
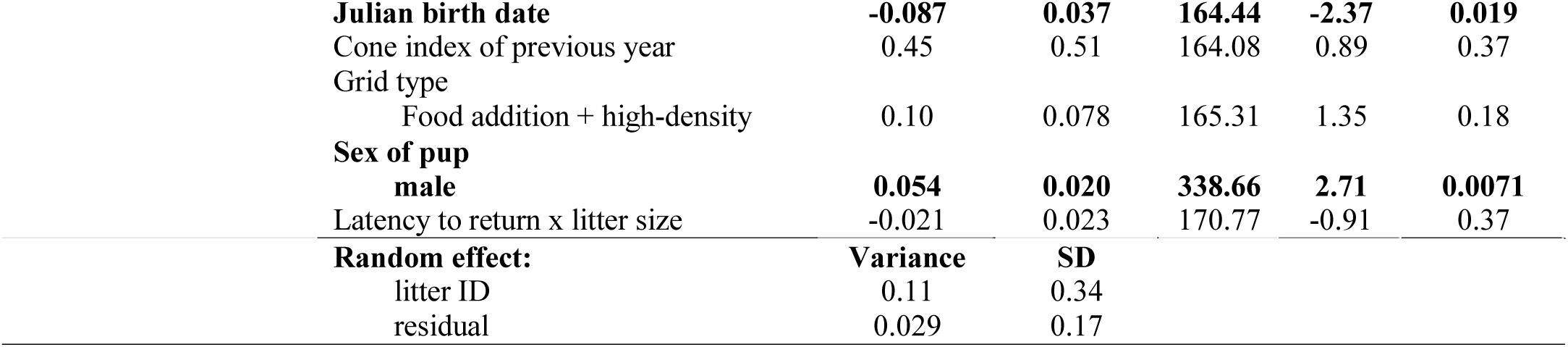
Results from our linear mixed-effects models on the relationship between growth rate and maternal attentiveness. We ran two distinct models for observations from the (a) first and (b) second nest entries (at ∼1 day post-parturition and ∼25 days post-parturition, respectively). We standardized litter size by grid type-year each combination and standardized all other continuous variables (latency to return to pups and cone indexes) across all data. Bold font indicates statistical significance of *p* < 0.05.

**a) Latency observations at first nest entry (pups ~1 day old):**
| Response Variable | Fixed Effect | b | SE | df | t | p-value |
| --- | --- | --- | --- | --- | --- | --- |
| **Growth rate (g/day)** |  |  |  |  |  |  |
| n = 524 individuals | Intercept | 1.15 | 0.94 | 180.78 | 1.23 | 0.22 |
| n = 193 litters | Latency to return to pups | -0.036 | 0.024 | 259.71 | -1.49 | 0.14 |
| n = 167 mothers | **Litter size** | **-0.14** | **0.025** | **192.53** | **-5.87** | **$p < 0.0001$** |
|  | Parity of mother |  |  |  |  |  |
|  | primiparous | -0.14 | 0.10 | 181.42 | -0.86 | 0.39 |
|  | Year |  |  |  |  |  |
|  | 2009 | 0.075 | 0.61 | 179.87 | 0.12 | 0.90 |
|  | 2016 | 0.46 | 0.94 | 180.72 | 0.49 | 0.63 |
|  | 2017 | 0.22 | 0.77 | 180.78 | 0.29 | 0.77 |
|  | **Julian birth date** | **-0.10** | **0.033** | **180.18** | **-3.07** | **0.0025** |
|  | Cone index of previous year | 0.34 | 0.72 | 180.62 | 0.72 | 0.47 |
|  | Grid type |  |  |  |  |  |
|  | Food addition + high-density | 0.081 | 0.071 | 180.97 | 1.14 | 0.26 |
|  | **Sex of pup** |  |  |  |  |  |
|  | male | **0.040** | **0.018** | **370.67** | **2.22** | **0.027** |
|  | **Latency to return x litter size** | **-0.051** | **0.023** | **196.22** | **-2.22** | **0.028** |
|  | **Random effect:** | **Variance** | **SD** |  |  |  |
|  | litter ID | 0.10 | 0.32 |  |  |  |
|  | residual | 0.027 | 0.16 |  |  |  |

**b) Latency observations at second nest entry (pups ~25 days old):**
| Response Variable | Fixed Effect | b | SE | df | t | p-value |
| --- | --- | --- | --- | --- | --- | --- |
| **Growth rate (g/day)** |  |  |  |  |  |  |
| n = 480 pups | Intercept | 0.82 | 1.00 | 164.21 | 0.83 | 0.41 |
| n = 178 litters | **Latency to return to pups** | **-0.06** | **0.028** | **160.96** | **-2.19** | **0.030** |
| n = 152 mothers | **Litter size** | **-0.11** | **0.024** | **175.09** | **-4.60** | **$p < 0.0001$** |
|  | Parity of mother |  |  |  |  |  |
|  | primiparous | -0.061 | 0.11 | 164.15 | -0.57 | 0.57 |
|  | Year |  |  |  |  |  |
|  | 2009 | 0.32 | 0.65 | 163.47 | 0.49 | 0.62 |
|  | 2016 | 0.81 | 1.00 | 164.14 | 0.81 | 0.42 |
|  | 2017 | 0.45 | 0.82 | 164.23 | 0.56 | 0.58 |

|  |  |  |  |  |  |
| --- | --- | --- | --- | --- | --- |
| **Julian birth date** | **-0.087** | **0.037** | **164.44** | **-2.37** | **0.019** |
| Cone index of previous year | 0.45 | 0.51 | 164.08 | 0.89 | 0.37 |
| Grid type |  |  |  |  |  |
| Food addition + high-density | 0.10 | 0.078 | 165.31 | 1.35 | 0.18 |
| **Sex of pup** |  |  |  |  |  |
| **male** | **0.054** | **0.020** | **338.66** | **2.71** | **0.0071** |
| Latency to return x litter size | -0.021 | 0.023 | 170.77 | -0.91 | 0.37 |
| **Random effect:** | **Variance** | **SD** |  |  |  |
| litter ID | 0.11 | 0.34 |  |  |  |
| residual | 0.029 | 0.17 |  |  |  |

As soon as lactation was detected through trapping, we used VHF radio telemetry to locate the nests where VHF collars were put on the mother. We will refer to this as the “first nest entry” (n = 292 litters from 167 females; mean pup age ± SD: 2.75 ± 3.33 days old). We estimated parturition date based on mass of pups, as well as palpation and lactation history of the mother (McAdam et al. 2007). When pups were ∼25 days old, we repeated this process for what we will refer to as the “second nest entry” (n = 227 litters from 152 females; mean pup age ± SD: 25.8 ± 2.47 days old). Forty-eight females were observed across two years.

At each nest entry, we removed pups from the nest to weigh (to nearest 0.1 g), mark, and sex individuals. During the second nest entry, we additionally assigned unique colored disk combinations and unique alphanumeric stamped ear tags (National Band and Tag Company, Newport, KY, USA) to each pup for identification after emergence. In a sample of nests from 2016, the average time pups spent out of the nest during the first nest entry was 11:56 min:sec [range: 06:19-34:50] (n = 65 litters) and the average time pups spent out of the nest during the second nest entry was 38:32 [range: 14:47-01:31:17] (n = 64 litters). In general linear models, we found no relationship between the time pups spent out of the nest for processing and the latency to return to pups (ß = −35.37, SE = 110.44, t_185_ = −0.32, *p* = 0.75) or the number of pups in a litter (ß = −75.27, SE = 144.23, t_185_ = −0.52, *p* = 0.60). This suggests that the time pups spent outside of the nest for processing or the number of pups in a litter did not influence the latency for a female to return to her nest. Data on time pups spent out of the nest were not collected in other years, but the above data should be representative of all years as data collection protocols were uniform across all years.

After processing the litter, we replaced all pups in the original nest. While the pups are being processed, there is high variation in maternal behavior. Some mothers explore the empty nest and stay nearby or approach the researchers. Some vocalize during the entire process and never approach. Mothers may even enter the nest before the researcher has left the nest tree. Alternatively, some mothers will move further away from the researchers or immediately leave and only return once the researchers have left (Westrick, personal observation). Not every litter was observed for both first and second nest entries because some of the litters did not survive from the first to second nest entry and some first nest entries were missed.

After each nest entry, we performed focal behavioral observations on mothers. An observer (n = 31 different observers) moved >5 m away from the nest tree and watched the mother’s behavior for 7 minutes following replacement of pups in the nest to record the time pups were replaced in the nest, the time mother returned to the nest, and the time mother began moving the pups. Observers were blind to the previous return latencies of the focal squirrel and were blind to the specific growth rates of pups. Because the observers processed the litter prior to behavioral observations, it was impossible to keep them blind to the litter size. Five observations were at underground nests, with the remaining in trees. We determined latency to return to pups as the time between pup replacement in their original nest and the mother’s return to the nest and censored any observations where the mother did not return within 7 minutes (n = 319 censored observations). Mothers typically moved their pups to a different nest immediately following their return to the nest after our intrusion as indicated by a strong relationship between the uncensored latency to return and latency to begin moving pups (linear model: adjusted R^2^ = 0.81, ß = 0.87, SE = 0.032, t = 27.43, *p* < 0.0001). Among trials where the mother returned, 83% of mothers moved their pups within 7 minutes. By measuring latency to return, our goal was to capture individual variation in how motivated a mother was to retrieve her pups to move them to a safer nest following a nest intrusion.

While many studies in the lab measure the time to retrieve all pups, the spatial scale at which wild female red squirrels move their offspring makes this problematic. Females move their pups individually to a new nest that is meters to tens of meters away. As a result, variation in the length of time between initiation and completion of moving pups is likely to be caused mostly by the distance between nests and the number of pups to be moved rather than by maternal motivation. Recording the latency to return to her pups following a standardized nest disturbance allows us to quickly capture the responsivity of a mother to her pups needs in a wild animal.

### Offspring measurements: growth rate and survival

Growth between the two nest entries is approximately linear (McAdam and Boutin 2003b), so we calculated growth rate (g/day) of pups (n = 671 pups) as the change in mass from first to second nest entry divided by number of days between nest entries. We monitored survival of juveniles (n = 870 juveniles from 251 litters) for the remainder of the year and following spring. We recorded survival to autumn of the birth year as a binary measure (alive or dead on August 15^th^). As part of our long-term data collection, we censused the entire study population yearly to confirm territory ownership by August 15^th^ and again by May 15^th^ (McAdam et al. 2007). Survival to August 15^th^ captures a key life-history event. In this population, caching of spruce cones typically begins mid-August and ends in September (Fletcher et al. 2010). Territory ownership before this period allows individuals to take advantage of that year’s cone crop by providing them with a physical space to cache cones (cones must be cached in a midden for the seed to remain a viable food source (Streubel 1968)). Because offspring disperse from their natal territory around 70-80 days old to compete for their own territory (Nice et al. 1956; McAdam et al. 2007), we limited survival data to litters born ≥ 70 days prior to August 15^th^. Because squirrels are diurnal and their activity (territorial defense behavior and presence) is conspicuous, we were able to completely enumerate all squirrels inhabiting the study areas through a combination of repeated live trapping and behavioral observations. We are confident our survival observations are not influenced by dispersal outside our study area because, in this population, the apparent survival of juveniles born on the edge of the study areas is not significantly different from those born in the core (Kerr et al. 2007).

From our biannual population censuses and behavioral observations, we tracked lifetime reproductive success (LRS) of mothers. We defined LRS as the number of pups borne over the entire lifetime of a dam that survived to recruit into the breeding population (i.e., alive for more than 199 days or roughly to the spring following their year of birth). To accurately calculate LRS, we only included mothers with known birth years before 2011 (n = 45 females) to ensure we captured the number of pups produced over their entire lifespan. We excluded mothers that died of unnatural causes.

All work was conducted under animal ethics approvals from Michigan State University (AUF#04/08-046-00), University of Guelph (AUP#09R006), and University of Michigan (PRO00005866).

### Statistical analyses

We conducted all statistical analyses in R version 3.4.3 (R Core Team, 2016). To estimate within-individual repeatability of maternal attentiveness, we used the R package ‘rptR’ version 0.9.21 (Stoffel et al. 2017). In our linear mixed effects model for repeatability, we included squirrel identity (ID) as a random intercept term, no fixed effects, and used parametric bootstrapping (n = 1000) to estimate the confidence interval.

In our models to assess how maternal nest attentiveness affected offspring growth, we included the following predictors: return latency, number of pups in litter, parity of mother (first time mother or not), Julian parturition date of the litter, cone index of the previous autumn, sex of pup, birth year (as a factor), and study area (control or high-density). We used the R package ‘lme4’ version 1.1-19 (Bates et al. 2015) to fit linear mixed-effects models and estimated P-values using the R package ‘lmerTest’ version 3.0-1 (Kuznetsova et al. 2016). To detect any collinearity in the predictors included in our model, we used R package ‘car’ version 3.0-2 (Fox and Weisberg 2011) to assess the variance inflation factors. GVIF^(1/(2xDF))^ for all predictors was < 3, except cone index of the previous autumn (GVIF^(1/(2xDF))^ = 11) which is colinear with birth year. We decided to still include spruce cone abundance (cone index: LaMontagne et al. 2005) in these models as it is a major influence on offspring survival and growth rate in this study system (McAdam and Boutin 2003a; McAdam and Boutin 2003b; Boutin et al. 2006; Dantzer et al. 2013) and we wanted to control for its influence on these traits. We included birth year to control for any additional year effects, e.g. variation in predator abundance or weather effects. Cone index of the previous year is also predictive of conspecific density which also influences growth rate (Dantzer et al. 2013). To assess if maternal nest attentiveness behavior could mitigate effects of increasing litter size on offspring growth rates, we included the interaction between return latency and litter size. We standardized pup growth rates, litter size, and birthdate within each grid-year combination and standardized all other continuous variables across all data (latency to return and cone index). Since multiple pups were measured per litter, we included litter ID as a random effect.

To model the relationship between maternal care and offspring survival, a binary value for offspring survival to autumn was predicted by the following fixed effects: return latency, pup growth rate, sex of pup, parity of mother (first time mother or not), Julian birth date, cone index of birth year, cone index of previous year, and study area (control or high-density). We standardized pup growth rate across grid-year combinations and standardized all other continuous variables (latency to return to pups and cone indexes) across all data. Again, we assessed variance inflation factors and found GVIF^(1/(2xDF))^ for all predictors was < 2.

Due to the count nature of LRS and the high variance of LRS relative to the mean, we used a negative binomial generalized linear model to estimate the relationship between maternal attentiveness and LRS. For each squirrel, we averaged latency to return to pups across all observations of that individual. In addition to latency to return to the pups, the model included fixed effects for lifespan (in days) since lifespan is highly correlated with LRS (McAdam et al. 2007). While mothers who experience a spruce cone mast in their lifetime have higher LRS on average (Haines et al. unpublished), in our dataset experiencing a mast year is highly correlated with lifespan (Pearson’s correlation R: 0.87, t = 11.78, df = 43, *p* < 0.00001), therefore we left this out of the model. We also fit the same model reducing the dataset to only observed return latencies, excluding any censored data, as a comparison. To fit these models, we used the R package ‘MASS’ version 7.3-51.1 (Venables and Ripley 2002). We standardized all continuous fixed effects to allow for comparison of effect size. GVIF^(1/(2xDF))^ for all predictors was < 2.

We ran all models for growth rate and survival with observations from the two nest entries separately due to the potential for different levels of maternal investment at different times in the breeding season. Specifically, squirrels born earlier in the year generally are more likely to survive until the following year so mothers that lose their litter earlier in the season (e.g. right after birth) have the potential to successfully breed again (McAdam et al. 2007; Williams et al. 2014), whereas mothers that lose their litter later in the season (e.g. a month after birth of the first litter) may not have the same potential for a successful second litter in a non-mast year. Additionally, newborn pups are hairless and more dependent on their mother for temperature regulation than ∼25-day old pups with fur. Consequently, we might expect behavior observations at the two nest entries to vary due to this difference in maternal investment and pup developmental stage, thus the measurements at the two nest entries may not be equivalent. Additionally, due to natural litter failures and missed observations, not every litter was observed at both the first and second nest entries.

## Results

### Variation and repeatability in maternal nest attentiveness

Mothers varied in the time it took them to return to the nest following our temporary removal of their pups (median: 385 s, CI = [333, 420]). Observations were censored at 7 minutes (50% of observations) and the latency to return ranged from 0 s to the maximum observation of 420 s. During 16% of observations, mothers returned within 2 minutes. Using Kaplan-Meier survival curves, the median latency to return was slightly faster during the second nest entry compared to the first nest entry, but survival curves do not significantly differ (nest entry 1 median = 407 s; nest entry 2 median = 342 s; difference between curves: χ^2^ = 0.1, p = 0.7).

In our models for within-individual repeatability of latency to return to pups, we found mothers were consistent across observations of maternal nest attentiveness behavior whether we included censored observations (R = 0.22, SE = 0.053, CI = [0.11, 0.32], p = 0.000013), or excluded them (R = 0.36, SE = 0.09, CI = [0.17, 0.52], p = 0.00072).

### Maternal nest attentiveness and pup growth rate

We found that the apparent cost for an individual pup of being in a litter with many siblings, in terms of a reduced growth rate, was lessened by having a highly attentive mother. As predicted by life history theory, in both our models of growth rate, pups from larger litters grew more slowly than pups in small litters (nest 1: ß = −0.14, SE = 0.025, t_192.53_ = −5.87, *p* < 0.00001; nest 2: ß = −0.11, SE = 0.024, t_175.09_ = −4.60, *p* < 0.00001). However, pups in large litters with mothers that returned soon after pup replacement (i.e., more attentive mothers) grew faster than those in large litters with mothers that took longer to return to the nest, particularly early in pup development (latency x litter size interaction - nest entry 1: ß = −0.051, SE = 0.023, t_196.22_ = - 2.22, *p* = 0.028; nest entry 2: ß = −0.021, SE = 0.023, t_170.77_ = −0.91, *p* = 0.37; Figure 1; Table 1).

**Figure 1.**
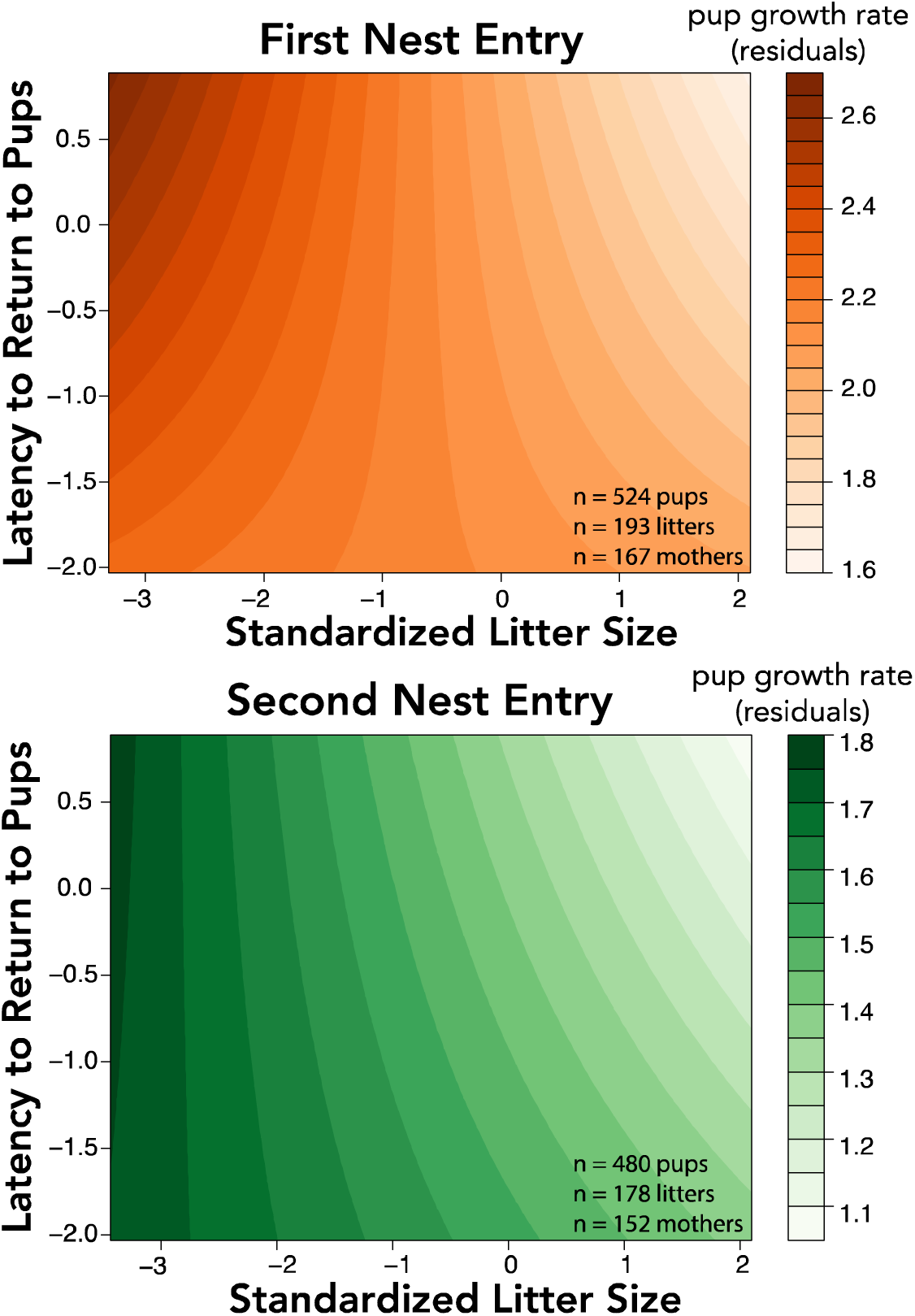
Pups from larger litters typically grew slower than pups from smaller litters. This relationship was mitigated by highly attentive mothers, at both the (A) first nest entry and (B) second nest entry, though the relationship was not statistically significant for the second nest entry (Table 1). Pups in large litters grow faster if their mother was highly attentive for latency to return to the nest at the first nest entry. More saturated, darker colors indicate a faster growth rate. Partial residuals of growth rate are plotted. Both litter size and latency to return to pups are standardized.

In both models for the first and second nest entry, male pups grew faster than female pups (nest entry 1: ß = 0.040, SE = 0.018, t_370.67_ = 2.22, *p* = 0.027; nest entry 2: ß = 0.054, SE = 0.020, t_338.66_ = 2.71, *p* = 0.0071; Table 1) and pups born later in the year also grew faster than pups born earlier (nest entry 1: ß = −0.10, SE = 0.033 t_180.18_ = −3.07, *p* = 0.0025; nest entry 2: ß = −0.087, SE = 0.037, t_164.44_ = −2.37, *p* = 0.019, Table 1).

### Maternal nest attentiveness and survival

Faster growing pups in both models were more likely to survive to autumn (nest entry 1: ß = 0.24, SE = 0.11, z = 2.12, *p* = 0.034; nest entry 2: ß = 0.29, SE = 0.12, z = 2.44, *p* = 0.015; Table 2). There was no further effect of maternal attentiveness once the direct effect of growth was accounted for (nest entry 1: ß = −0.11, SE = 0.11, z = −1.00, *p* = 0.32; nest entry 2: ß = −0.12, SE = 0.11, z = −1.05, *p* = 0.29; Table 2). Overall, females were more likely than males to survive until autumn (nest entry 1: ß = 0.71, SE = 0.22, z = −3.30, *p* < 0.0001; nest entry 2: ß = −0.54, SE = 0.23, z = −2.34, *p* = 0.019; Table 2). Pups born earlier in the year (nest entry 1: ß = −0.27, SE = 0.12; z = −2.17, *p* = 0.00098; nest 2: ß = −0.19, SE = 0.13, z = −1.47, *p* = 0.14; Table 2) and pups born during years where there was high autumn spruce cone production were more likely to survive (nest entry 1: ß = 0.25, SE = 0.11, z = 2.21, *p* = 0.027; nest 2: ß = 0.36, SE = 0.14, z = 2.64, *p* = 0.0084; Table 2).

**Table 2.**
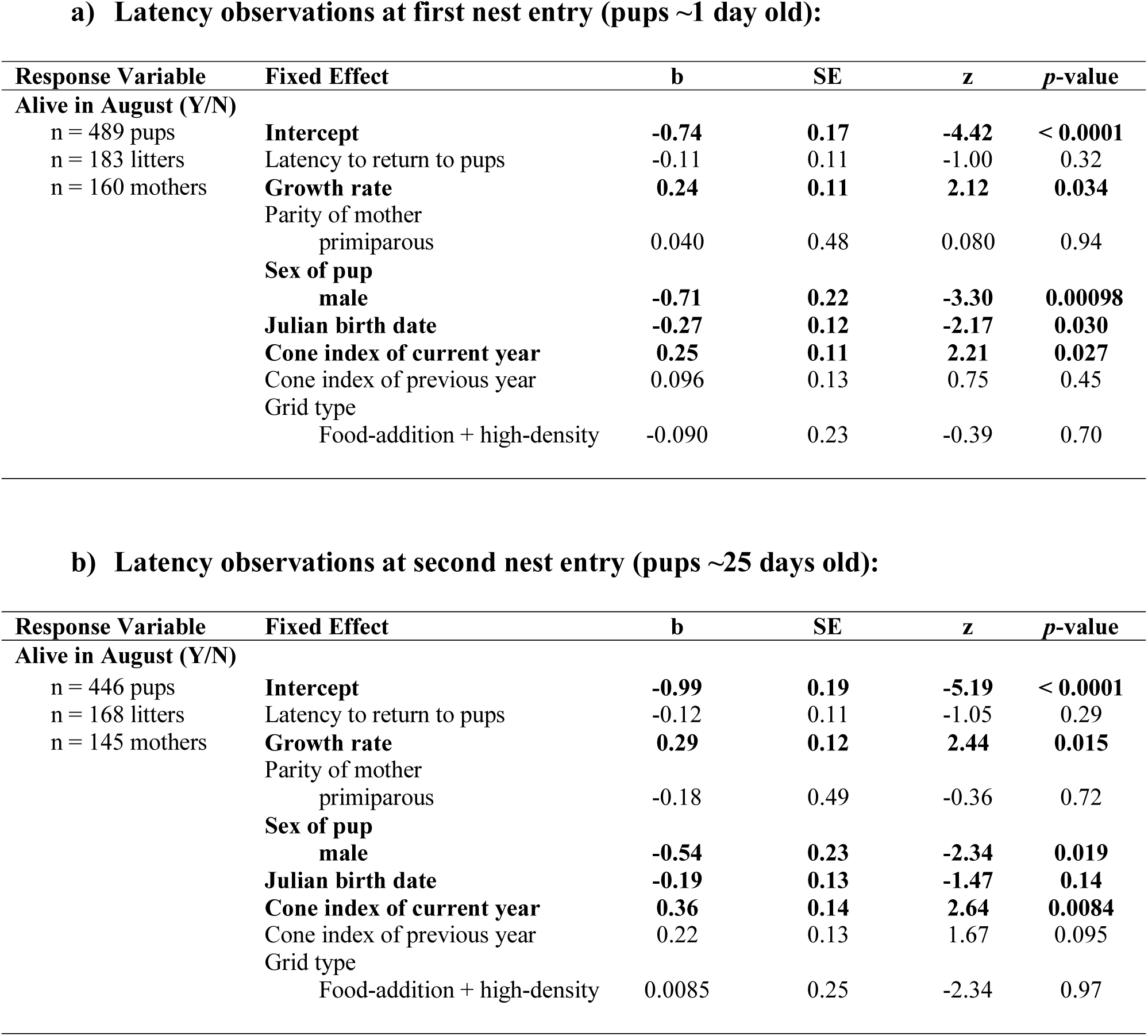
Results from our binomial linear models on the relationship between survival to autumn and maternal attentiveness. We ran two distinct models for observations from the (a) first and (b) second nest entries (at ∼1 day post-parturition and ∼25 days post-parturition, respectively). We standardized growth rate (g/day) and Julian birth date within grid-year combination. We standardized all other continuous variables (latency to return to pups and cone indexes) across all data. Bold font indicates statistical significance of *p* < 0.05.

**a) Latency observations at first nest entry (pups ~1 day old):**
| Response Variable | Fixed Effect | b | SE | z | p-value |
| --- | --- | --- | --- | --- | --- |
| Alive in August (Y/N)<br>n = 489 pups<br>n = 183 litters<br>n = 160 mothers | **Intercept** | **-0.74** | **0.17** | **-4.42** | **< 0.0001** |
|  | Latency to return to pups | -0.11 | 0.11 | -1.00 | 0.32 |
|  | **Growth rate** | **0.24** | **0.11** | **2.12** | **0.034** |
|  | Parity of mother |  |  |  |  |
|  | primiparous | 0.040 | 0.48 | 0.080 | 0.94 |
|  | **Sex of pup** |  |  |  |  |
|  | male | **-0.71** | **0.22** | **-3.30** | **0.00098** |
|  | **Julian birth date** | **-0.27** | **0.12** | **-2.17** | **0.030** |
|  | **Cone index of current year** | **0.25** | **0.11** | **2.21** | **0.027** |
|  | Cone index of previous year | 0.096 | 0.13 | 0.75 | 0.45 |
|  | Grid type |  |  |  |  |
|  | Food-addition + high-density | -0.090 | 0.23 | -0.39 | 0.70 |

**b) Latency observations at second nest entry (pups ~25 days old):**
| Response Variable | Fixed Effect | b | SE | z | p-value |
| --- | --- | --- | --- | --- | --- |
| Alive in August (Y/N)<br>n = 446 pups<br>n = 168 litters<br>n = 145 mothers | **Intercept** | **-0.99** | **0.19** | **-5.19** | **< 0.0001** |
|  | Latency to return to pups | -0.12 | 0.11 | -1.05 | 0.29 |
|  | **Growth rate** | **0.29** | **0.12** | **2.44** | **0.015** |
|  | Parity of mother |  |  |  |  |
|  | primiparous | -0.18 | 0.49 | -0.36 | 0.72 |
|  | **Sex of pup** |  |  |  |  |
|  | male | **-0.54** | **0.23** | **-2.34** | **0.019** |
|  | **Julian birth date** | **-0.19** | **0.13** | **-1.47** | **0.14** |
|  | **Cone index of current year** | **0.36** | **0.14** | **2.64** | **0.0084** |
|  | Cone index of previous year | 0.22 | 0.13 | 1.67 | 0.095 |
|  | Grid type |  |  |  |  |
|  | Food-addition + high-density | 0.0085 | 0.25 | -2.34 | 0.97 |

### Maternal nest attentiveness and lifetime reproductive success

Mothers who on average returned to their pups faster had slightly more pups recruit into the population during their lifetime (ß = −0.35, SE = 0.19, z = −1.89, *p* = 0.059; Figure 2; Table 3a). While this relationship is uncertain and therefore not statistically significant, the effect size is quite large. On average, mothers who returned immediately after pups were replaced in the nest had ∼1 more pup recruit than mothers who returned at the end of the 7 min observation period. If we only include observed maternal behavior data (n = 32 females, no censored data), this relationship becomes stronger (ß = −0.40, SE = 0.17, z = −2.31, *p* = 0.021; Table3b). Females who lived longer had higher LRS (ß = 0.56, SE = 0.16, z = 3.40, *p* = 0.00068; Table 3a).

**Figure 2.**
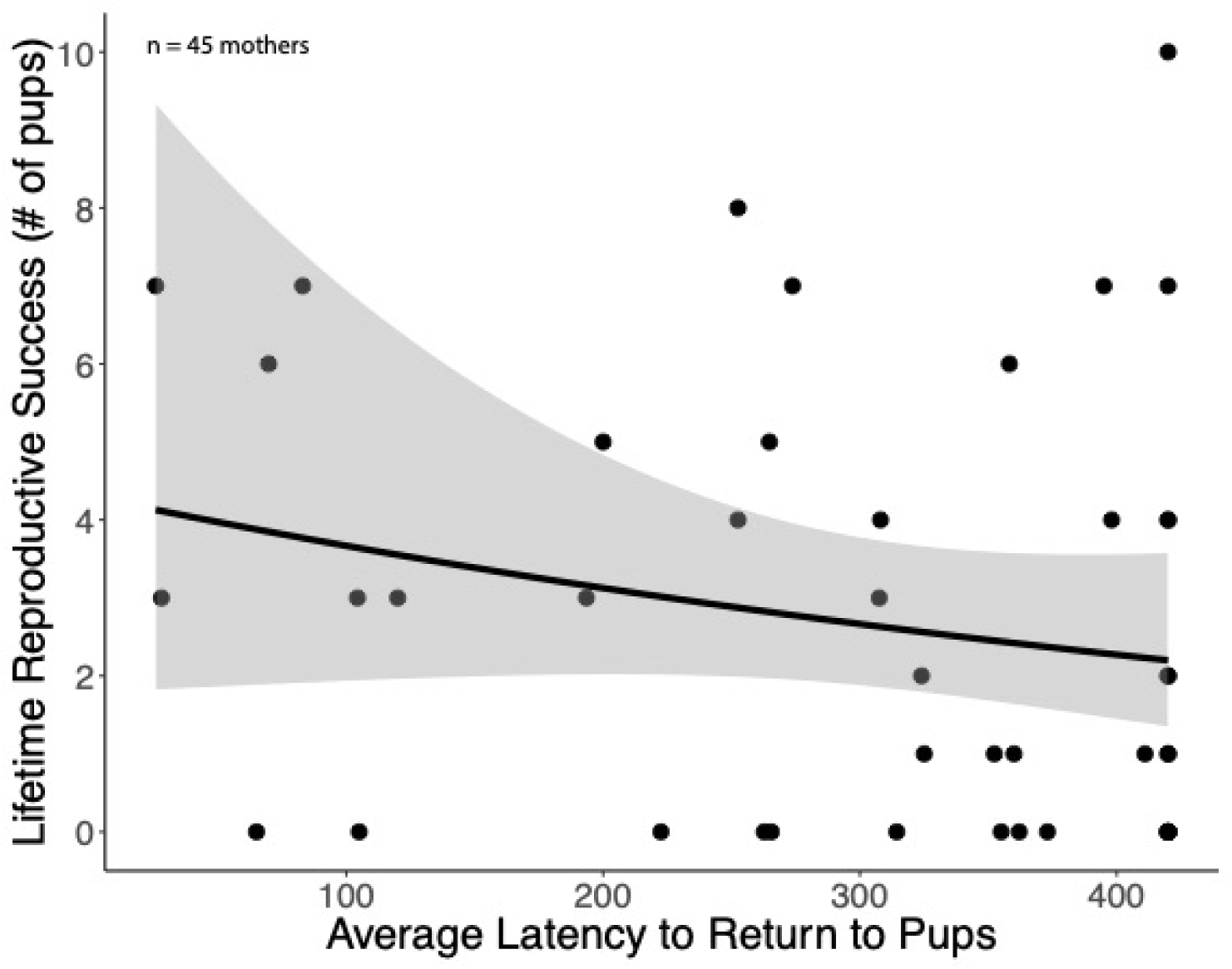
Highly attentive mothers have more pups survive to recruit into the breeding population than less attentive mothers.

**Table 3.** Results from our negative binomial linear models on the relationship between lifetime reproductive success of females and maternal motivation. We defined lifetime reproductive success as the number of pups born surviving over winter to recruit into the breeding population. For each squirrel, we averaged latency to return to pups across all observations of that individual, including censored observations as 420 s for (a) and removing any censored observations for (b). We standardized both continuous fixed effects. We included squirrels with known birth years prior to 2011 (range: 2004 to 2010). Bold font indicates statistical significance of *p* < 0.05.

**a) Latency to return to pups (censored at 7 min)**
| Response Variable | Fixed Effect | b | SE | z | p-value |
| --- | --- | --- | --- | --- | --- |
| **Lifetime reproductive success (# of pups)**<br>n = 45 mothers |  |  |  |  |  |
|  | **Intercept** | **0.87** | **0.17** | **5.23** | **< 0.0001** |
|  | Average latency to return to pups | -0.35 | 0.19 | -1.89 | 0.059 |
|  | **Lifespan (days)** | **0.56** | **0.16** | **3.40** | **0.00068** |

**b) Latency to return to pups (within 7 minutes)**
| Response Variable | Fixed Effect | b | SE | z | p-value |
| --- | --- | --- | --- | --- | --- |
| **Lifetime reproductive success (# of pups)**<br>n = 32 mothers |  |  |  |  |  |
|  | **Intercept** | **0.97** | **0.17** | **5.57** | **< 0.0001** |
|  | Average latency to return to pups | -0.40 | 0.17 | -2.31 | 0.021 |
|  | **Lifespan (days)** | **0.44** | **0.16** | **2.68** | **0.0074** |

## Discussion

We found that maternal nest attentiveness (measured as latency to return to pups) following a nest intrusion is a repeatable behavior among female red squirrels, which suggests individuals exhibit maternal styles with some mothers being more attentive to the pups and other mothers adopting a more laissez-faire approach. The repeatability of this maternal behavior is near the average repeatability of other behavior studies (R = 0.37; Bell et al. 2009), but lower than the repeatability of other behavioral traits in red squirrels (docility R = 0.41, aggression R = 0.44, activity R = 0.51; Taylor et al. 2012). Research in wild deer mice (*Peromyscus maniculatus*) also demonstrates low, but significant, repeatability of multiple maternal behaviors with a high degree of seasonal plasticity (Stewart and McAdam 2014). We are limited in assessing the plasticity of return latency behavior in red squirrels as our dataset does not include observations during a mast year, a potential driver of a plastic change in behavior.

Variation in maternal behavior in red squirrels has important consequences for offspring. Female red squirrels exhibiting higher levels of maternal nest attentiveness lessened the fundamental life history trade-off between offspring quantity and quality. Red squirrel mothers that were highly attentive at the first nest entry were capable of producing faster growing pups, compared to less attentive mothers, by mitigating the trade-off between litter size and pup growth rates. This amelioration of the negative impact that siblings can have on the growth of each offspring in the litter could be one way that maternal behavior alters offspring lifetime fitness trajectories (Klug and Bonsall 2014). For example, offspring that grow up in large litters may grow nearly as fast as offspring in smaller litters if they have a highly attentive mother. Mothers could potentially decrease the amount of time pups spend in a vulnerable life stage by increasing growth rate or, alternatively, promote slower development in poor environmental conditions (Klug and Bonsall 2014). Studies in other species have found that parents can alter their investment in the quantity or quality of offspring according to environmental conditions. For example, in quacking frogs (*Crinia georgiana*), mothers face a trade-off between egg size and number and can use variable egg provisioning to influence survival of offspring in good and poor-quality environments (Dziminski and Roberts 2006). Similarly, in birds, habitat elevation is an important factor influencing whether parents invest in quantity or quality of offspring (Badyaev and Ghalambor 2001). Our study suggests red squirrels use maternal behavior as one mechanism to adjust the trade-off between offspring size and number, but the ecological cause of variation in maternal behavior is yet to be determined. Mothers may invest heavily in offspring when density is high to increase growth or reduce investment when the benefits no longer exceed the costs during years when growth is less predictive of success, such as mast years (Humphries and Boutin 2000; McAdam and Boutin 2003a; Dantzer et al. 2013). Indeed, we see higher rates of litter loss in mast years potentially suggesting less maternal investment and care (Haines et al. 2018).

In our study population of red squirrels, faster early life growth rate is associated with an increased probability of pup survival into adulthood, especially when population density is high (McAdam and Boutin 2003a; Dantzer et al. 2013; Hendrix et al. 2019). Because growth rate is predicted by maternal behavior, including these two measurements as predictors in the same model may mask the indirect impact of maternal care on survival. These results suggest growth rate may be the mechanism by which maternal behavior increases survival of pups, but we have no evidence to suggest that maternal attentiveness has any further effect on offspring survival beyond that which is mediated through the increased growth rate of her pups. Additionally, we found that, over their lifetime, mothers that exhibit a more attentive maternal style have more offspring that recruit into the breeding population. Latency to return to pups and subsequently move them to a new nest following a nest disturbance may affect offspring growth and survival through a variety of pathways. Lactating female red squirrels are known to move their pups between nests on their territory as the ambient temperature fluctuates to maintain an optimum temperature for offspring growth (Guillemette et al. 2009). Since latency to return to pups is highly correlated with latency to move pups (see Methods), highly attentive mothers may be better able to move offspring from one nest to another that puts offspring in the optimal thermal environment that maximizes growth. Highly attentive mothers may also increase offspring growth or survival by reducing flea load on pups by reducing their risk of death due to infanticide (Haines et al. 2018) or predation (Studd et al. 2014), or by moving them to nests with lower infestations of ectoparasites (Gooderham and Schulte-Hostedde 2011; Duma 2013). It is also likely that maternal attentiveness, as we measured it here, does not have any direct effects on growth and survival but is simply representative of a suite of maternal behaviors representing maternal style.

Despite finding that highly attentive mothers had higher lifetime reproductive success, we found substantial individual variation in maternal nest attentiveness. This begs the question of why is there variation in this highly beneficial behavior? Although, we did not explicitly compare the costs and benefits assumed in parental investment theory, there are likely costs experienced by highly attentive mothers. For example, if the nest was intruded upon by predators, highly attentive mothers that quickly return to the nest could face the cost of potentially being preyed upon themselves. Additionally, there are likely substantial energetic costs associated with moving pups to a new nest; on average one ∼25-day old pup weighs ∼18% of the body mass of an adult female. We have not yet documented the costs of maternal attentiveness but there are three possible explanations for why there is substantial individual-variation in maternal attentiveness despite the clear fitness benefits we documented in this study. First, high maternal nest attentiveness could be exhibited by high quality mothers who can afford higher investment in current reproduction, and variation we see in maternal behavior is due to limitations on the mother and current environmental conditions, rather than fitness consequences (van Noordwijk and de Jong 1986). For example, highly attentive mothers that produce fast growing offspring with higher survival may be those with larger amounts of cached food.

Secondly, it is entirely possible that the survival costs to females of increased attentiveness are underrepresented in our data due to the ‘invisible fraction’, or individuals that do not survive to express this behavior (Grafen 1988; Hadfield 2008). We are not able to measure maternal attentiveness on all squirrels and many squirrels die prior to even reproducing so we are unable to collect data on them. We are more likely to have sampled older individuals for maternal attentiveness which means the survival costs to females with increased attentiveness are likely to be underrepresented in our data. Essentially, a cost associated with increased attentiveness could have already been paid prior to us being able to measure attentiveness.

Lastly, there may also be years when the fitness benefits associated with maternal attentiveness are reduced. Red squirrels in Yukon experience large fluctuations in their major food resource and there is substantial inter-annual variation in directional selection on offspring growth rate (McAdam and Boutin 2003a; Dantzer et al. 2013). In some years, there is strong positive selection favoring fast growth whereas it is reduced or non-significant in other years. Consequently, it may be an unreliable strategy for mothers to invest in faster growing offspring if it can result in a high energetic cost with little fitness benefit (McAdam and Boutin 2003a; Dantzer et al. 2013). This should result in balancing selection on maternal attentiveness where less attentive mothers with slow growing pups have an advantage in years when fast growth is not under positive selection, thereby maintaining individual-variation in maternal attentiveness. While we are limited in testing these predictions with our current dataset, we have already observed balancing selection for other repeatable behavioral traits in female red squirrels such as the aggression and activity of mothers (Taylor et al. 2014). Given that individual variation in maternal behavior has also been documented in other species (e.g. Rachlow and Bowyer 1994; Fairbanks 1996; Dahle and Swenson 2003; Ringsby et al. 2009), it is possible that the variable fitness benefits associated with maternal care depending upon ecological conditions is likely to maintain variation in this important behavior that is closely linked to fitness.

## Acknowledgements

We thank Agnes MacDonald and her family for long-term access to her trapline, and Champagne and Aishihik First Nations for allowing us to conduct our work within their traditional territory. We thank all volunteers, field assistants, and graduate students for their assistance in data collection. This work was supported by American Society of Mammalogists to SEW; University of Michigan to SEW and BD; National Science Foundation (IOS-1749627 to BD, DEB-0515849 to AGM); and Natural Sciences and Engineering Research Council to SB, AGM, and JEL. This is publication XX of the Kluane Red Squirrel Project.

